## Supplemental Figures for "Automated single-cell proteomics providing sufficient proteome depth to study complex biology beyond cell type classifications"

### Title

Steven A. Carr

Broad Institute of MIT and Harvard
415 Main Street
02142 Cambridge
MA, USA.

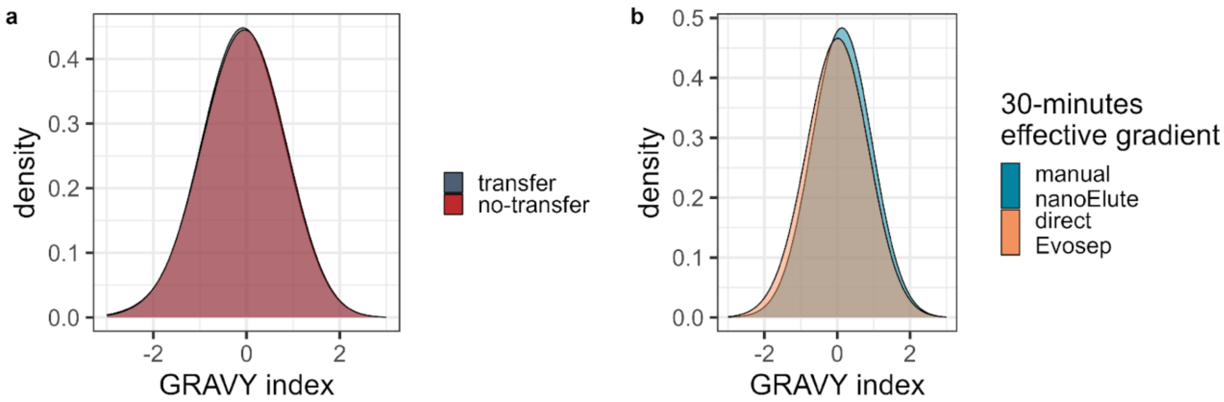


Supplemental Fig. 1: GRAVY index of peptides identified in (a) ddaPASEF with manual transfer (gray; n = 5) and automated transfer via centrifugation (red; n = 5) or in (b) diaPASEF with 30-minutes effective gradients on the nanoElute with manual transfer to a HPLC vial (dark green; n = 25) or automated transfer to the Evotip and acquisition with 40SPD (orange; n = 25).

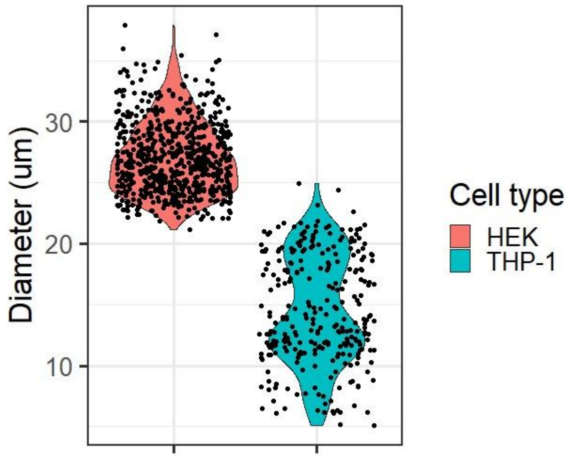


Supplemental Fig. 2: Density distribution of dispensed HEK-293T (n = 672) and THP-1 (n = 288)

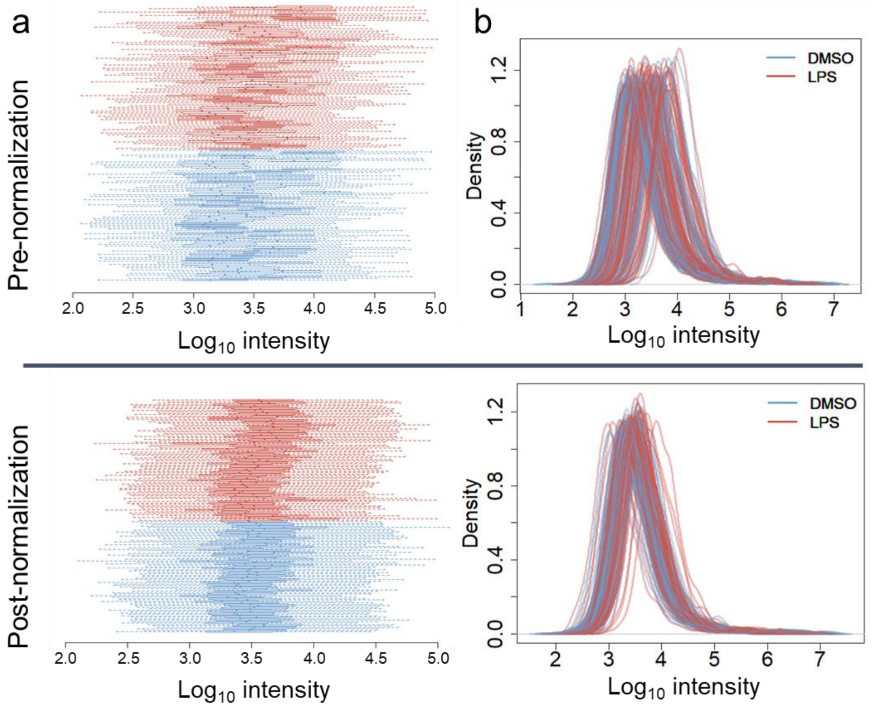


cell diameters.

Supplemental Fig. 3: Boxplots (a) and profile plots (b) of $\log_{10}$-transformed protein intensities before (top) and after (bottom) normalization using SCnorm. Each boxplot/density represents the distribution of proteins from one cell. Cells are colored by LPS treatment.

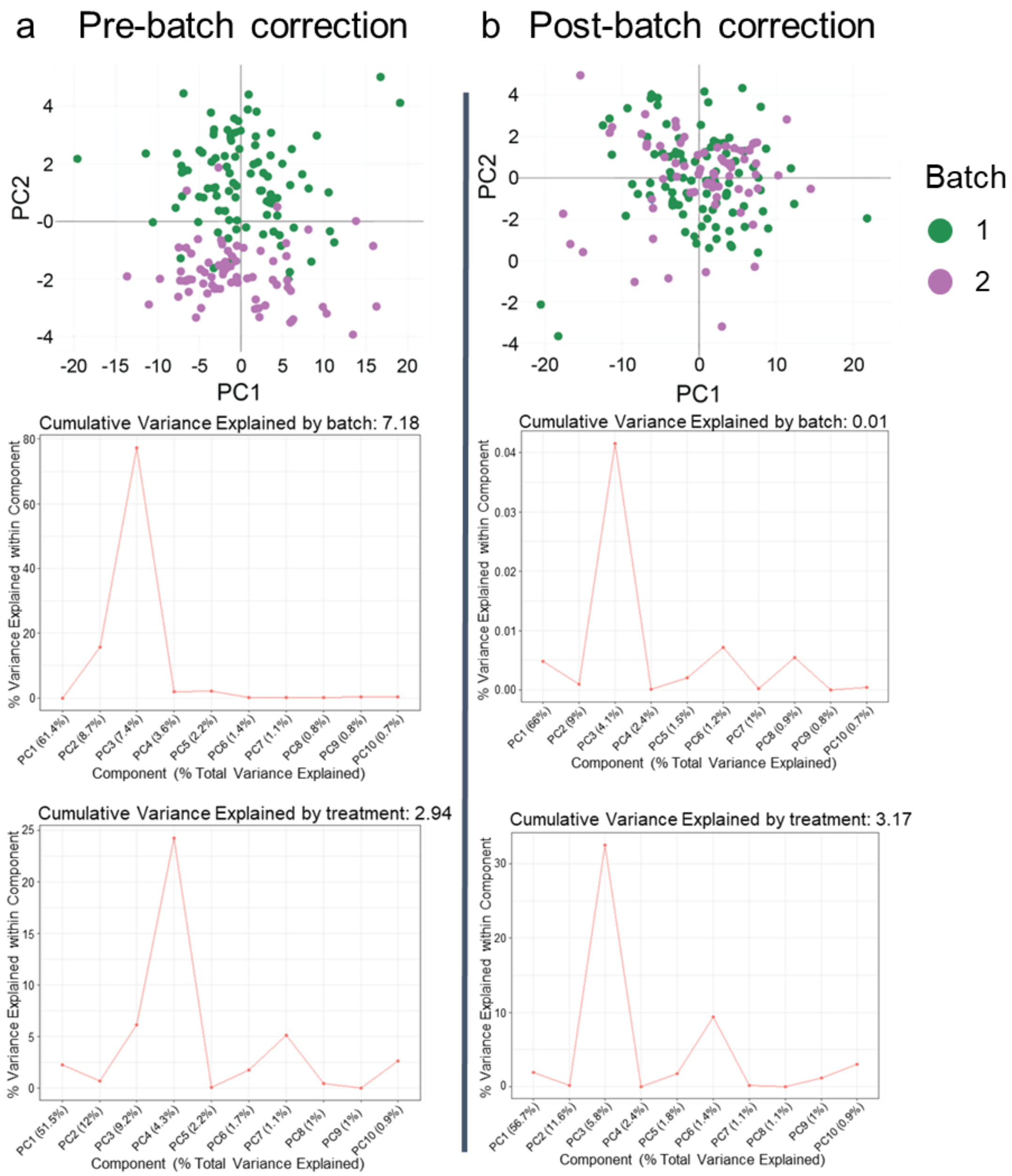


Supplemental Fig. 4: PCA and PC regression of LPS treated (n = 84) with DMSO control (n = 77) THP-1 cells (a) pre- and (b) post batch correction. PC regressions indicates cumulative variance explained by batch or group (= treatment). Colors indicate experimental batches.

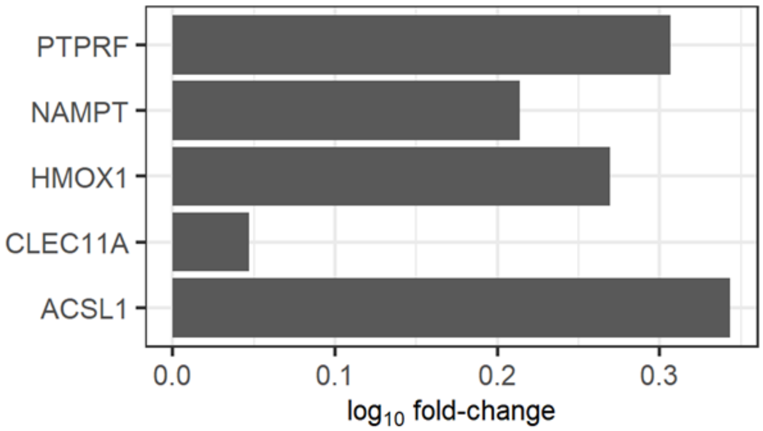


Supplemental Fig. 5: $Log_{10}$ fold-change of previously published LPS response proteins in THP-1 cells (n = 161) upon 12 hours LPS treatment.
